# DNA methylation-based forensic age estimation in human bone

**DOI:** 10.1101/801647

**Authors:** Shyamalika Gopalan, Jonathan Gaige, Brenna M. Henn

**Affiliations:** Department of Ecology and Evolution, Stony Brook University, Stony Brook, NY 11794, USA; Department of Anthropology, University of California, Davis, CA 95616, USA; UC Davis Genome Center, University of California, Davis, CA 95616, USA

## Abstract

DNA methylation is an epigenetic modification of cytosine nucleotides that represents a promising suite of aging markers with broad potential applications. In particular, determining an individual’s age from their skeletal remains is an enduring problem in the field of forensic anthropology, and one that epigenetic markers are particularly well-suited to address. However, all DNA methylation-based age prediction methods published so far focus on tissues other than bone. While high accuracy has been achieved for saliva, blood and sperm, which are easily accessible in living individuals, the highly tissue-specific nature of DNA methylation patterns means that age prediction models trained on these particular tissues may not be directly applicable to other tissues. Bone is a prime target for the development of DNA methylation-based forensic identification tools as skeletal remains are often recoverable for years post-mortem, and well after soft tissues have decomposed. In this study, we generate genome-wide DNA methylation data from 32 individual bone samples. We analyze this new dataset alongside published data from 133 additional bone donors, both living and deceased. We perform an epigenome-wide association study on this combined dataset to identify 108 sites of DNA methylation that show a significant relationship with age (FDR < 0.05). We also develop an age-prediction model using lasso regression that produces highly accurate estimates of age from bone spanning an age range of 49-112 years. Our study demonstrates that DNA methylation levels at specific CpG sites can serve as powerful markers of aging, and can yield more accurate predictions of chronological age in human adults than morphometric markers.

## Introduction

Determining a person’s age from skeletal and dental remains has been an integral component of building a biological profile for several decades of forensic investigations. Classic methods rely on morphological features of development and functional decline, such a tooth eruption, suture closure, and bone density (Cunha et al., 2009; Franklin, 2010). However, these approaches suffer from difficulties in implementation and standardization (Cunha et al., 2009; Franklin, 2010). One such issue is that there are few morphological features that can be used to precisely estimate the age of adults, as most developmental processes that can distinguish between juveniles of different ages cease by adulthood. Thus, different skeletal features are analyzed for this purpose in different age classes, but the overall accuracy of morphology-based estimates is typically low for adults (Cunha et al., 2009; Franklin, 2010).

Molecular-based methods of age estimation represent a promising alternative to those based on morphometrics. One early approach was based on the degree of amino-acid racemization of bones and teeth, which produces highly accurate estimates of age (Meissner and Ritz-Timme, 2010; Ohtani and Yamamoto, 2010). However, this method still suffers from practical limitations, such as difficulties in standardizing the procedure, and the need for specialized equipment and technical expertise to isolate the tissue of interest and conduct protein fractionation (Meissner and Ritz-Timme, 2010; Ohtani and Yamamoto, 2010).

More recently, advances in the study of DNA methylation have driven its popularity in the field of forensic sciences as a general molecular approach to estimating individual age that can be applied in various contexts. DNA methylation is a chemical modification to the primary DNA sequence; in mammals, it occurs predominantly on cytosine nucleotides that are in a cytosine-guanine dinucleotide context (a CpG site). It has been shown that, at certain CpG sites, the average level of DNA methylation changes measurably throughout life in a manner that makes them potentially useful markers of human aging. This has led to a profusion of methods being developed for tissues such as sperm, blood, saliva, and teeth which produce highly accurate estimates based on DNA methylation levels from relatively few CpG sites, measured using widely used, often commercially available, assay methods (Hong et al., 2017; Lee et al., 2015; Vidaki et al., 2017; Yi et al., 2014; Zbieć-Piekarska et al., 2015b). However, there does not yet exist a DNA methylation-based method for estimating age from human bone, a tissue type that is highly relevant to the field of forensic sciences. Methods developed for other tissues likely cannot be applied directly to bone due to the highly tissue-specific nature of age-related DNA methylation patterns (Dmitrijeva et al., 2018; Hannum et al., 2013; Lee et al., 2015; Maegawa et al., 2010; Slieker et al., 2018).

In this study, we analyze bone-derived DNA methylation data from hundreds of thousands of CpG sites across the genome in a large sample of human adults to identify CpG sites that change significantly with age. We then use a subset of these sites to develop a highly accurate prediction model for individuals in an age class where morphological methods can be problematic. Furthermore, we show that our model exhibits higher accuracy and relies on fewer CpG sites compared to a previously published model that was developed for use on multiple tissues.

## Methods

### Description of datasets

Three previously analyzed and published datasets were derived from bone biopsies of living donors, assayed on the Illumina Human Methylation 450k array, and previously analyzed (Table 1) (Horvath et al., 2015; Reppe et al., 2017). Additional data were generated for this study from bone-derived DNA from deceased donors from both forensic and preserved specimens and assayed on the Illumina Human Methylation EPIC array (Table 1).

**Table 1.** Description of datasets analyzed.

| **Dataset (GEO accession)** | **N samples** | **N females** | **Age range** | **Sample type** | **Methylation array** | **Citation** |
| --- | --- | --- | --- | --- | --- | --- |
| Osteoporosis study | 84 | 84 | 49-86 | Living donor | 450k | (Reppe et al., 2017) |
| Multi-tissue study (GSE64490) | 48 | 46 | 49-104 | Living donor | 450k | (Horvath et al., 2015) |
| Supercentenarian from multi-tissue study (GSE64491) | 1 (plus 3 replicates) | 1 | 112 | Deceased donor | 450k | (Horvath et al., 2015) |
| Preserved specimens | 19 | 7 | 60-97 | Deceased donor | EPIC | This study |
| Forensic specimens | 13 | 4 | 18-95 | Deceased donor | EPIC | This study |

### Forensic specimens

Forensic specimens were collected from the Forensic Anthropology Center at Texas State (FACTS) in San Marcos, Texas. Donated specimens are typically allowed to decompose naturally in an outdoor environment on the premises for 2-3 years, after which the bones are cleaned with a mild detergent and housed in their storage facilities. For this study, trabecular bone from femurs was collected based on a previously published protocol for minimally destructive sampling for skeletal remains (Gibbon et al., 2009). Briefly, the outside of the bone was cleaned between the medial and lateral condyles with 95% ethanol. The femur was then held vertically while a small hole was drilled with a Dremel 200 rotary tool and a 1/8” diameter bit. After rotating the fully inserted drill bit and withdrawing it, the femur was inverted over a collection tube to collect the bone powder. Between 0.1 and 1g (0.67g average) of bone powder was recovered for these samples.

### Preserved specimens

Preserved specimens were collected from the Department of Anatomical Sciences Stony Brook University. Bodies donated to this facility are treated with chemical preservatives and used in medical and dental anatomy courses. For each sample, a Dremel 200 rotary tool with a 1” cutting disc was used to extract a piece of tissue that contained the petrous portion of the temporal bone, which has been previously shown to be among the densest skeletal elements in the body and typically contains relatively uncontaminated endogenous DNA in even ancient samples (Pinhasi et al., 2015). This piece was then pre-digested in a solution of TE buffer and proteinase K at 55°C for up to 5 days, or until all soft tissue was dissolved, leaving behind only bone. This piece was then subsampled into smaller pieces and ground to a fine powder using a combination of a mortar and pestle and an IKA tube mill. Between 0.27 and 1.13g (0.54g average) of bone power per sample was extracted.

### DNA extraction

Bone samples underwent a pre-digestion step prior to DNA extraction using a standard phenol-chloroform method. The powder was incubated in a solution of 250 uL of N-Laurylsarcosine and 30uL proteinase K in 10mL EDTA at 55°C on a nutating mixer for two days to dissolve the mineral structure of the bone. If powder was not completely dissolved after two days, an additional 5mL of EDTA was added and the sample incubated for a further two days. One volume of phenol-chloroform was then added to the dissolved bone solution, shaken at room temperature for one minute, and centrifuged for 20 minutes at 3,500 rpm. The aqueous phase was transferred to a fresh tube where one volume (the same amount as EDTA and phenol-chloroform) of chloroform was added. The sample was again shaken for one minute and centrifuged for 20 minutes. The aqueous phase was then transferred to an Amicon Ultra-15 filter and centrifuged for up to 20 minutes, or until most of the liquid passed through. The filter was washed with 12mL of ultra-pure HPLC water and spun down until most of the liquid passed through. If necessary, this step was repeated until the wash-through liquid was clear.

### Methylation array data processing and quality control

DNA methylation data generated for this study was preprocessed from raw intensity files (IDATs) using the R package minfi (Aryee et al., 2014). The GEO deposited datasets were preprocessed from raw methylated and unmethylated counts using the R packages lumi and methylumi (Davis et al., 2015; Du et al., 2008). Raw data was not available for the osteoporosis study, so pre-processed and rescaled data (by beta mixture quantile dilation – BMIQ) provided by the author was used (Reppe et al., 2017). All raw data were first filtered for poorly performing samples and CpG sites. Samples were flagged if their median methylated or unmethylated levels were low (if log2 of the count was less than 10.5), if more than 10% of measured probes exceeded a detection p-value threshold of 1%, or if the sex predicted by X and Y chromosome methylation levels did not match the reported sex, as per standard DNA methylation array processing procedures (Aryee et al., 2014). All forensic samples failed these quality control measures. Probes that were among previously identified cross-reactive probes were removed (Chen et al., 2013; Pidsley et al., 2016). Colour correction, background correction, and functional normalization of IDAT data was conducted with the ‘preprocessFunnorm’ function in minfi (Aryee et al., 2014). Colour correction, background correction, and shift and scaling normalization of signal count data was conducted with the ‘lumiMethyC’ and ‘lumiMethyN’ functions in lumi (Du et al., 2008). Rescaling of type 1 and 2 probes was conducted by BMIQ, implemented in the R package wateRmelon. All remaining probes that failed a detection p-value threshold of 5% were then set to NA and all CpG sites on sex chromosomes were also removed. Continuous beta values for each CpG site, which range from 0 (indicating that the site is completely unmethylated) to 1 (completely methylated), were extracted for subsequent analysis.

### Principal components analyses

We conducted a principal components analyses (PCA) on the merged dataset of samples that were successfully assayed (n = 155) in order to identify factors correlated with significant sources of variation across the datasets. PCAs were conducted with the ‘prcomp’ function in R.

### Epigenome-wide association study (EWAS)

An EWAS for age was performed using the R package CpGassoc on the combined dataset of 155 individuals and all CpG sites that were present on the 450k dataset and passed quality control. Dataset identity was included as a covariate to control for excessive genomic inflation in the EWAS.

### Horvath age prediction

The original Horvath age prediction algorithm was implemented by using the ‘agep’ function from the R package wateRmelon on the preprocessed and normalized data (Horvath, 2013; Pidsley et al., 2013). The optional normalization step in the Horvath workflow was not used.

### Developing age prediction models

We created a training set of data to develop an age prediction model by randomly sampling approximately 70% of individuals from each dataset, except for GSE64491, which contains four replicates of one individual; these were evenly divided into the test and training dataset. We then made minor label adjustments to ensure that the training set always contained the widest possible age range. We then used lasso regression, implemented in the R package glmnet (Friedman et al., 2010), to select predictors from only those CpG sites that were found to be significantly associated with age (FDR < 0.05) in the EWAS. We varied the regularization penalty parameter (λ – not to be confused with the genomic inflation factor) across a range to select two primary models: the best model had the lowest prediction error in the training dataset, and the other model had a prediction error within one standard error of the minimum in the training dataset.

### Gene region enrichment analysis

The 37 CpG sites comprising the best performing model were tested for enrichment of several possible categories, including gene and disease ontology terms, phenotypes, genes, and Molecular Signatures Database (MSigDB) terms using Genomic Regions Enrichment of Annotation Tool (GREAT) (McLean et al., 2010). A custom background set of all CpGs tested in the EWAS was used.

## Results

### Data quality

Genome-wide methylation data was successfully generated for all preserved specimens, but all forensic samples failed post-assay quality control measures. For the latter, both median methylated and unmethylated signals were too low to pass the threshold and all samples had a high proportion of unreliable probes (see *Methods*). Furthermore, while the median methylated signal was significantly lower than the median unmethylated signal across all samples (p = 9.2 × 10−6, paired sample t-test), this difference was more pronounced in the forensic samples (p = 4.1 × 10−14, t-test) (Figure 1). The forensic samples were excluded from all subsequent analyses.

**Figure 1.**
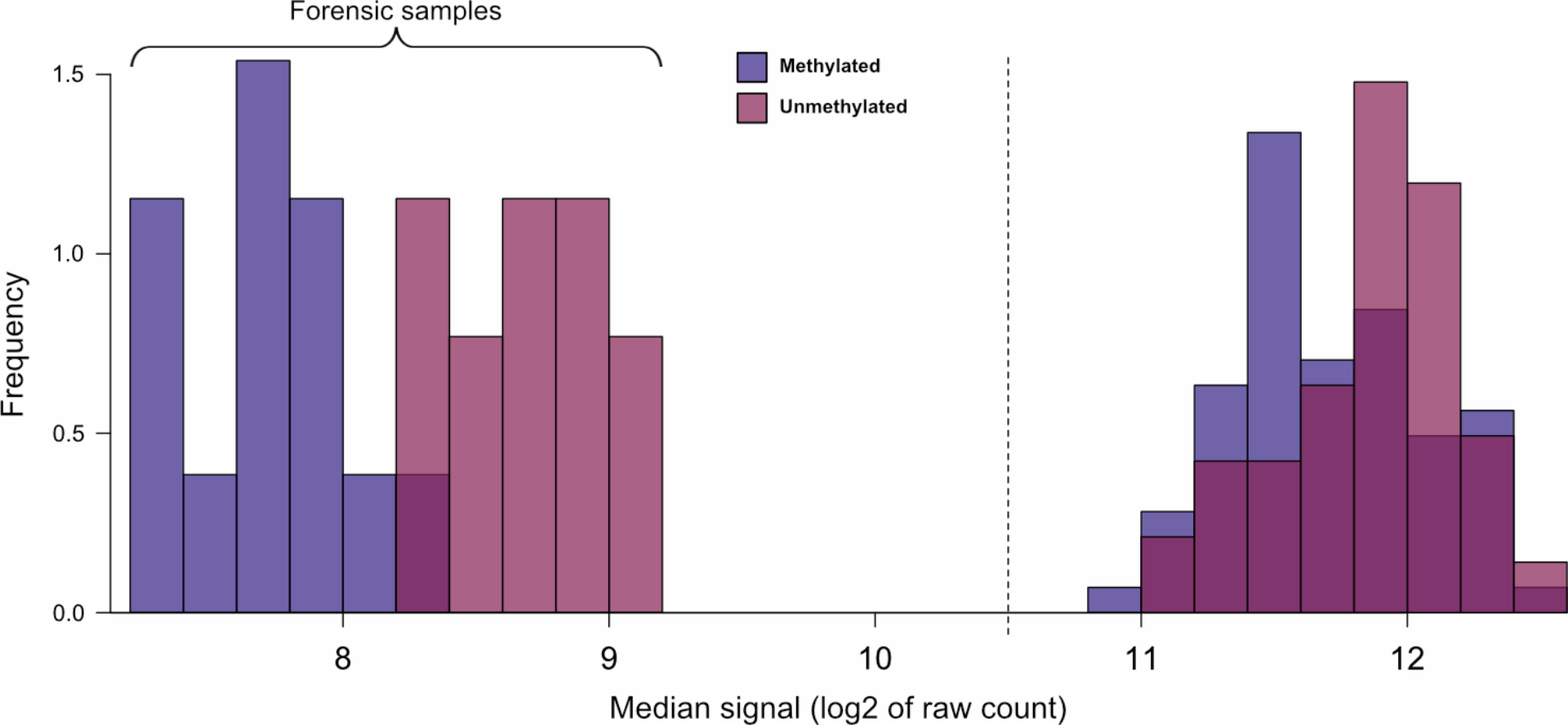
The distribution of median methylated (blue) and unmethylated (red) signal values in forensic samples (left) and all other samples (right). The threshold typically used for post-assay quality control is indicated with the dashed line (Aryee et al., 2014). All forensic samples in this study failed to meet this threshold and were thus excluded from subsequent analysis.

### Principal components analysis

A principal components analysis of the 155 successfully assayed samples showed that batch effects were the major drivers of the first two principal components, which together accounted for over 55% of the variation in the dataset (Figure 2).

**Figure 2.**
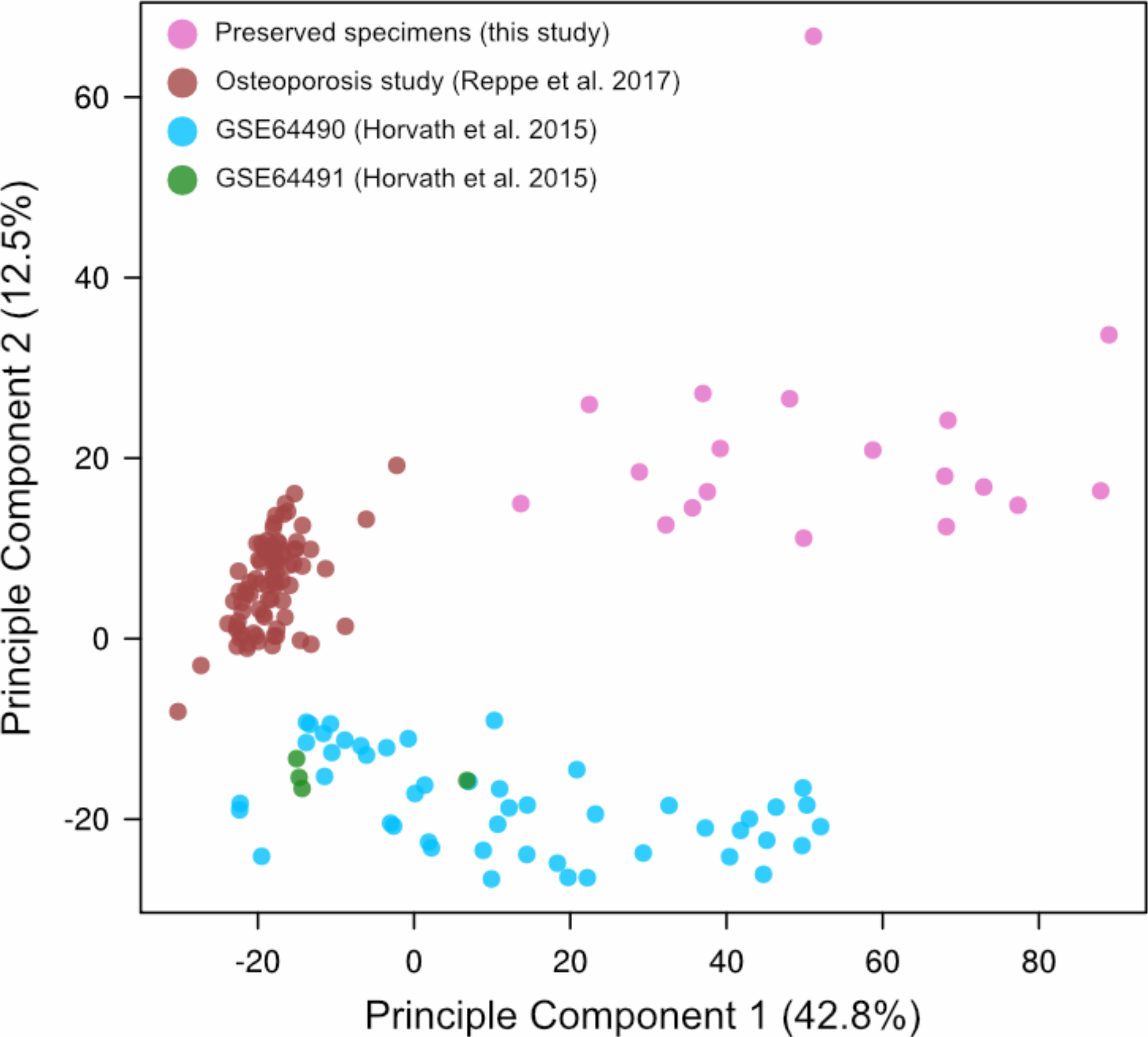
Scatterplot of the first two principal components of variation in the DNA methylation dataset.

### Epigenome-wide association study (EWAS)

We performed an EWAS on the merged DNA methylation dataset to identify CpG sites that were significantly associated with age in bone tissue (see *Methods*). When not correcting for dataset identity, we observed extreme genomic inflation, which is not unexpected given the substantial batch effects observed and the differences in age distributions across the three largest datasets (Figures 2–3; Table 1). When using dataset identity as a covariate in the EWAS, this inflation was greatly reduced, and 108 CpG sites were significantly associated with age (FDR < 0.05) (Figures 3–4).

**Figure 3.**
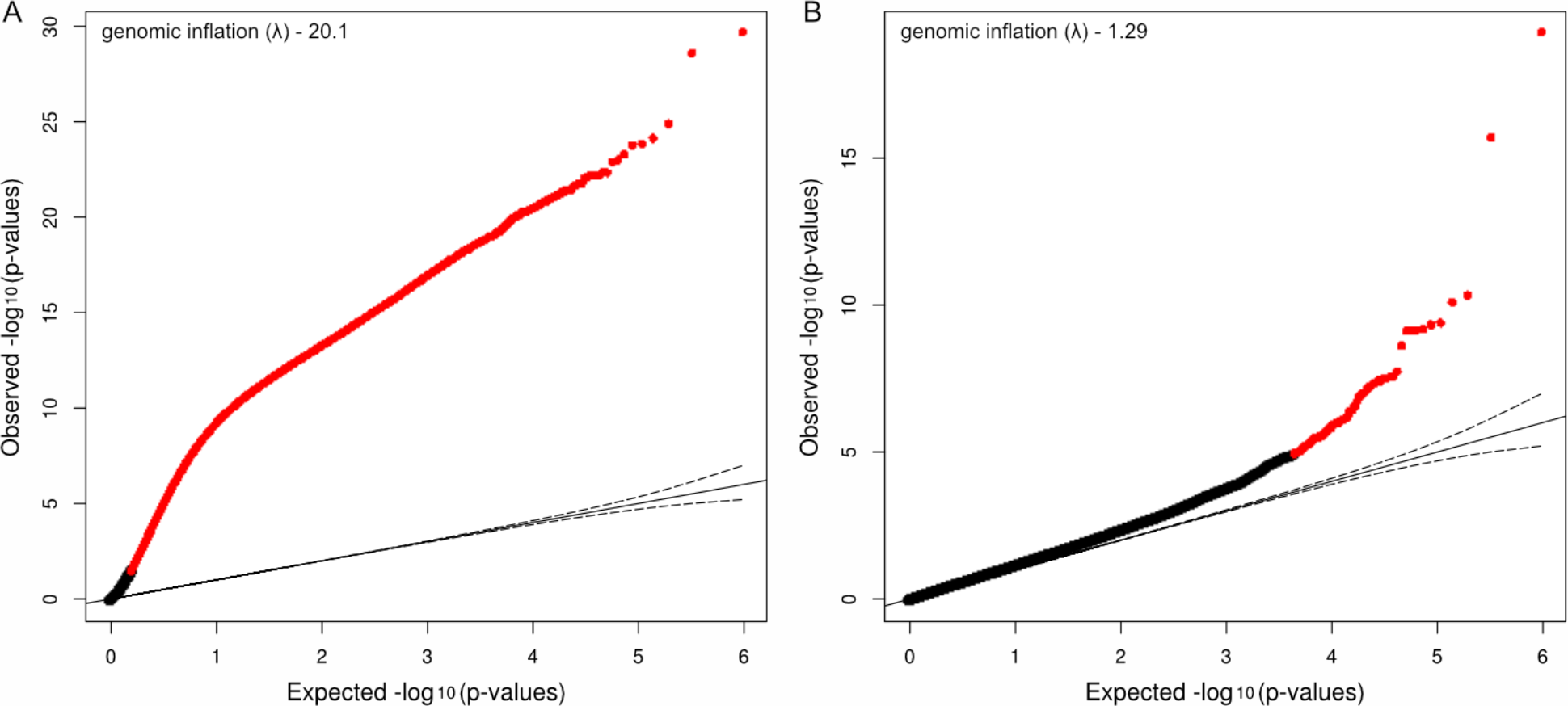
Quantile-quantile plot of expected p-values (under a uniform distribution) versus observed p-values in the EWAS. The points represent p-values for a tested CpG site. Red points indicate those that are found to be significantly associated with age (FDR < 0.05). A) When no covariates are used, an excessive number of CpG sites are found to be significant. B) The inclusion of dataset identity appears to correct this, likely by accounting for batch effects between sets of samples that were assayed separately.

**Figure 4.**
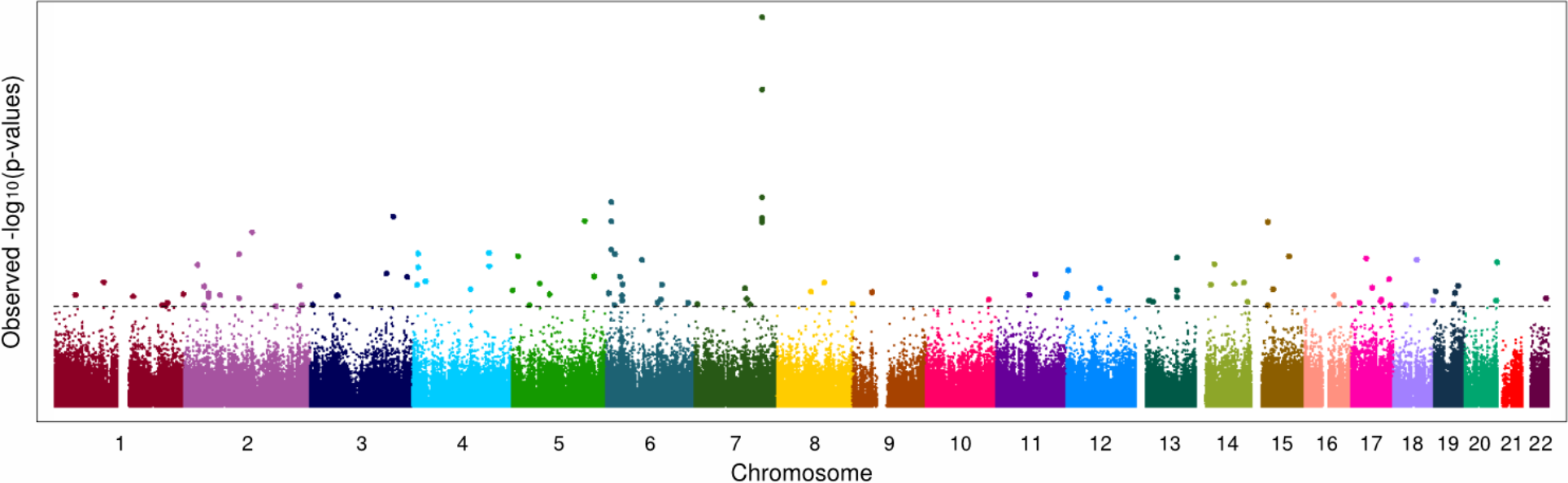
Manhattan plot depicting the location of all CpG sites tested in the EWAS where dataset identity was included as a covariate. The false discovery rate threshold of 0.05 is indicated by the horizontal dashed line. 108 CpG sites were found to exceed this threshold (larger dots).

### Developing and evaluating models for chronological age prediction for bone

We used the prediction model published by Horvath (2013), which leverages DNA methylation information from 353 CpG sites, to estimate the age of the individuals in the merged dataset. The model was trained on DNA methylation data derived from 13 different tissue types, including bone marrow, but not hard bone tissue (i.e. osteocytes and osteoblasts) (Horvath 2013). It was previously tested on the bone samples from the GSE64490 and GSE64491 datasets in Horvath et al. 2015 (see Figure 6 in that paper) and was consequently described as also being applicable to hard bone tissues, although no metric of accuracy was reported (Horvath et al 2015). We visually parsed 51 of the 52 data points in the GSE64490 and GSE64491 datasets using a plot digitizer (Rohatgi, 2018), and estimated the root mean squared error (RMSE) to be 8.1 years. Our application of the same model to these datasets, appears to have yielded more accurate results than the initial publication (5.9 and 6.3 years, respectively, and 6.0 across both datasets; Figure 5). The reasons for this are unclear, but could be attributed to differences in how the data were processed prior to running the age prediction algorithm (see *Methods* and *Discussion*).

**Figure 5.**
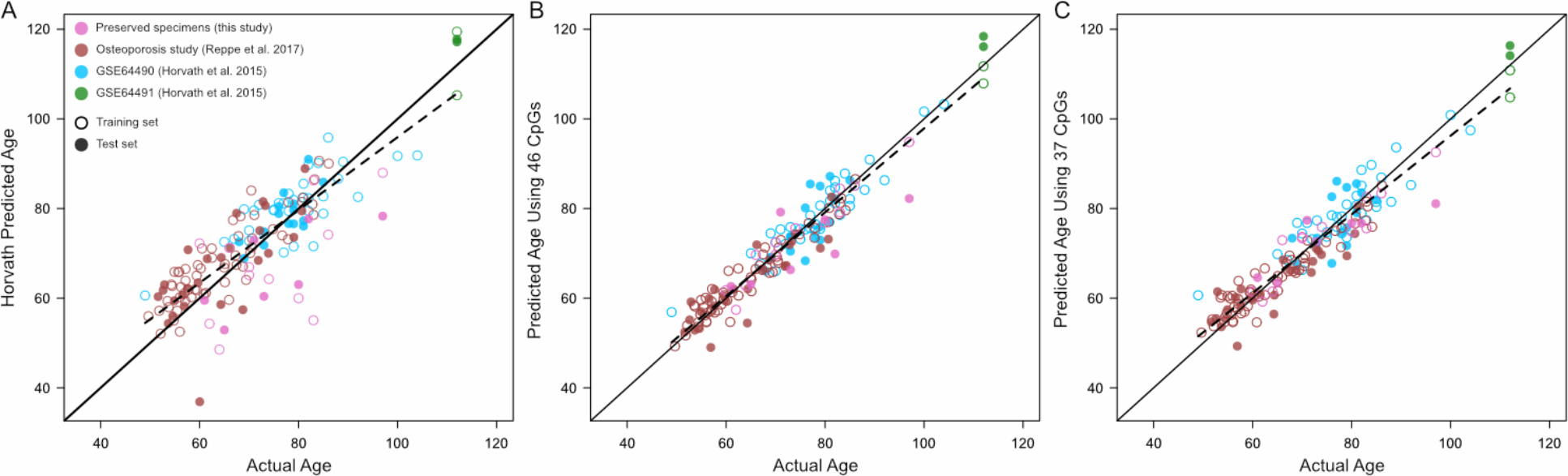
True chronological age versus estimates from three DNA methylation-based models. A) Age estimates from A) the Horvath model, based on DNA methylation data from 353 CpG sites, B) the 46-site model and C) the 37-site model. Outlined circles represent samples that were used to train the 46- and 37-site models, while solid circles represent test set samples.

We sought to determine if accurate estimates of individual age could be achieved for human bone using fewer than 353 sites. We split the individuals in the datasets into training and test groups and used lasso regression to select predictor variables from among the 108 age-associated CpGs identified in the EWAS (see *Methods*). The model that minimized the prediction error in the training data was based on information from 46 sites, while a second model exhibited comparably low error using information from only 37 CpG sites. Both models included two CpG sites that are also used in the Horvath model. These two new models were also used to generate estimates of age for all individuals in the dataset, and the RMSE between the true age and the prediction in the test dataset was calculated (Table 2). Both models were more accurate than the Horvath model, with the sparser model (based on 37 CpG sites) being slightly more accurate in the test samples overall (Figure 5; Table 2). This model is hereafter referred to as the ‘bone clock’, and additional details for the specific CpG sites that comprise it are given in Table 3.

**Table 2.** Root mean squared errors (RMSE) for each prediction model tested; the Horvath model, the 46-site model and the 37-site model. As the Horvath model was not trained on any of the individuals analyzed here, the RMSE for all samples is also reported. Of these three, the 37-site bone clock performs best in the test set of individuals.

| Model | Root mean squared error (years) |  |  |  |  |  |
| --- | --- | --- | --- | --- | --- | --- |
|  | All samples | Test samples |  |  |  |  |
|  |  | All test samples | Preserved samples (this study) | Osteoporosis study (Reppe et al. 2017) | GSE64490 (Horvath et al. 2015) | GSE64491 (Horvath et al. 2015) |
| *Horvath model* | 7.4 | 7.7 | 11.7 | 8.1 | 3.9 | 5.5 |
| *46 site model* | - | 5.1 | 8.4 | 4.2 | 4.6 | 5.4 |
| *37 site model* | - | 4.9 | 7.1 | 4.2 | 4.8 | 4.3 |

**Table 3.** Details of the 37 bone clock CpG sites. The names of all CpG sites that comprise our best model are reported here, along with their genomic positions (based on the hg19/GRCh37 reference), the gene(s) they are annotated to (if any), and their relative positions within those genes. The positions of these CpG sites are either within the transcriptional start site (TSS), gene body, exons, or untranslated regions (UTR). The two bone clock CpG sites that are also used by the Horvath clock are indicated by an asterisk.

| Name | Position | UCSC Gene Name | UCSC Gene Group |
| --- | --- | --- | --- |
| cg01974375 | 1:151298954 | PI4KB | TSS1500 |
| cg21748207 | 2:26395359 | FAM59B | TSS1500 |
| cg12206199 | 2:39187543 | LOC375196, LOC100271715 | TSS200, Body |
| cg20547295 | 2:69664869 | NFU1 | TSS200 |
| cg27213509 | 2:176947228 | EVX2 | Body |
| cg21545859 | 3:5068037 |  |  |
| cg03607117 | 3:53080440 | SFMBT1 | TSS1500 |
| cg07553761 | 3:160167977 | TRIM59 | TSS1500 |
| cg23995914 | 4:10459228 | ZNF518B | TSS200 |
| cg07171111 | 4:10462903 |  |  |
| cg01899437 | 4:24914441 | CCDC149 | 1<sup>st</sup> exon, 5'UTR |
| cg03133735 | 4:111562270 |  |  |
| cg03663715 | 5:72744904 | FOXD1 | TSS1500 |
| cg23500537 | 5:140419819 |  |  |
| cg13773570 | 6:6006707 | NRN1 | Body |
| cg16867657 | 6:11044877 | ELOVL2 | TSS1500 |
| cg24724428 | 6:11044888 | ELOVL2 | TSS1500 |
| cg00464814 | 6:16758889 | ATXN1 | 5'UTR |
| cg22736354* | 6:18122719 | NHLRC1 | 1<sup>st</sup> exon |
| cg10970124 | 6:31634602 | BAT4, CSNK2B | TSS1500, 5'UTR |
| cg13959344 | 6:32901642 |  |  |
| cg08541518 | 6:69942892 | BAI3 | Body |
| cg19509311 | 6:106429191 |  |  |
| cg00590036 | 6:158957433 | TMEM181 | TSS200 |
| cg07955995 | 7:130419159 | KLF14 | TSS1500 |
| cg24681895 | 8:145743681 | RECQL4, LRRC14 | TSS1500, 5'UTR |
| cg10217503 | 12:65153209 | GNS | 5'UTR, 1<sup>st</sup> exon |
| cg22171539 | 12:81107990 |  |  |
| cg11082362 | 14:36003181 | INSM2 | TSS200 |
| cg21801378* | 15:72612125 | BRUNOL6 | 1<sup>st</sup> exon |
| cg06279276 | 16:67184164 | B3GNT9 | Body |
| cg24668364 | 17:16284324 | UBB | TSS200 |
| cg22361181 | 17:40171740 | NKIRAS2 | TSS1500, 5'UTR |
| cg16969368 | 17:57642752 | DHX40 | TSS200 |
| cg17243289 | 18:45458021 | SMAD2 | TSS1500 |
| cg16665444 | 18:76828655 | ATP9B | TSS1500 |
| cg02997982 | 19:41082291 | SHKBP1, SPTBN4 | TSS1500, 3'UTR |

An enrichment analysis identified two significant categories as being overrepresented in these 37 bone clock CpG regions; the gene ELOVL2 was represented twice (a 948-fold enrichment relative to the background, p = 2.1 × 10−6), and genes bearing the motif ‘AAGCACA’ in their 3’ untranslated regions (UTRs) were represented 8 times (4.8-fold enrichment, p = 2. 1 × 10−4).

## Discussion

In this study, we present newly generated genome-wide DNA methylation data from adult human bones spanning a nearly 4 decade age range. We incorporate additional DNA methylation array data derived from living and deceased donors, resulting in a combined dataset that spans a total age range of over 6 decades, which we use to conduct the first comprehensive analysis of patterns of epigenetic aging in adult human bone tissue. We show that a previously published epigenetic age predictor generates reasonably accurate estimates of chronological age based on DNA methylation data from 353 CpG sites (RMSE 7.4) (Horvath 2013).

Surprisingly, we achieved more accurate estimates of chronological age using the Horvath algorithm on the GSE64490 and GSE64491 datasets than the original publication (RMSE of 6.0 years rather than ~8.1 years) (Horvath et al., 2015). This may be due to differences in processing the raw DNA methylation signal data. One major difference is our implementation of the beta mixture quantile dilation (BMIQ) normalization method, which corrects for technical differences between probes measured with the Type I and Type II Illumina chemistry, on all probes in the dataset (Teschendorff et al., 2013). By contrast, Horvath et al. uses a custom script to normalize data, which is based on the BMIQ method but uses a ‘gold standard’ reference panel of Type II probes (Horvath, 2013; Horvath et al., 2015). There may be additional differences in our processing pipelines beyond this normalization step, but it is nonetheless noteworthy that such significant variation in overall accuracy can result from such differences.

Relative to the best implementation of the Horvath model, I demonstrate that even more accurate estimates of chronological age can be generated for human bone from nearly 10 times fewer sites. There are two likely reasons for this improvement. The first is that the original Horvath model was only trained on CpG sites that were present on an earlier Illumina DNA methylation array, which assayed 27 thousand sites (Horvath 2013). This excluded any sites present on the more comprehensive 450k array, many of which were reported to be excellent potential predictors of age. For example, it was demonstrated that DNA methylation at the gene ELOVL2, and in particular the CpG site cg16867657, showed a strong and precise relationship with age (Garagnani et al., 2012). This association was confirmed in subsequent studies, which further supported its utility for estimating chronological age in humans with potential forensic applications (Hannum et al., 2013; Johansson et al., 2013; Naue et al., 2017; Spólnicka et al., 2018; Zbieć-Piekarska et al., 2015a). By training our predictor on data from age-associated CpG sites identified from 450k array data, we were able to achieve accurate results using fewer and, arguably, better markers of chronological age.

The second reason for improved accuracy is our focus on a single tissue rather than the Horvath model’s focus on multiple tissues. It is well known that genomic patterns of DNA methylation differ broadly across tissue types; in fact, DNA methylation is an important mechanism by which tissue identity is established during development (Lokk et al., 2014; Rakyan et al., 2008). That these tissue-specific differences intersect with changes in DNA methylation with age is also well established (Dmitrijeva et al., 2018; Hannum et al., 2013; Maegawa et al., 2010; Slieker et al., 2018). It is therefore unsurprising that it would be challenging to develop a predictor of age that is equally accurate on all human tissue types. In fact, the Horvath model has been used to argue that different tissues ‘age’ at different rates based on the consistent deviations from a linear model relating chronological age to the predicted age (Horvath et al., 2015, 2014).

It is important to note here a difference in the motivations behind the Horvath model and any model intended for forensic applications, including the one presented here. The Horvath model is primarily used as a measure of a conceptual ‘biological’, not chronological, age, which reflects individual functional capacity and overall health rather than simply the number of years lived. The 37 bone clock CpGs identified here may therefore be best suited for forensic applications, and not for estimating health status.

It is interesting to note that one of the significant results of the enrichment analysis was the gene ELOVL2. Two of 37 CpG sites selected by lasso regression, the aforementioned cg16867657 as well as cg23606718, were annotated to this gene. This supports the idea that the gene ELOVL2 is a uniquely useful predictor of chronological age, as it is significantly associated with age across multiple tissue types, including blood, saliva, brain, buccal cells, liver, fat, breast, kidney, lung, teeth, and now, bone, as well as across diverse populations (Bekaert et al., 2015; Giuliani et al., 2016; Gopalan et al., 2017; Hannum et al., 2013; Johansson et al., 2013; Slieker et al., 2018). However, it is still unclear if the specific relationship between DNA methylation level and age (i.e., the ‘slope’ and intercept’ of the linear model) is consistent across all these tissue types.

The second significant enrichment category identified here was the presence of the AAGCACA motif in the 3’ UTR. Genes that bear this motif are putative targets of negative regulation by the miR-218 microRNA, which carries the complementary sequence (Subramanian et al., 2005). It has been shown that miR-218 is upregulated during osteoblast differentiation, driving bone differentiation via a positive feedback loop involving the Wnt pathway (Hassan et al., 2012). Interestingly, while miR-218 has been shown to have tumor-suppressing activity in multiple tissue types, it also promotes metastasis of breast cancer to bone by mimicking this molecular signature of bone differentiation (Alajez et al., 2011; Hassan et al., 2012; Liu et al., 2018; Tatarano et al., 2011; Venkataraman et al., 2013; Yang et al., 2017). While it is unclear if the enrichment of miR-218 targets among the bone clock CpGs is meaningful, it is nevertheless tempting to speculate that it suggests a bone-specific characteristic of our predictive model.

A significant limitation of this study is the age range of individuals available for training the model. While the datasets altogether provided a cross-section of 63 years of adult human lifespan, it is not clear if our bone clock can accurately estimate the age of younger adults. In an attempt to improve age-association detection power by increasing the sample size and the age range surveyed, we also assayed 13 forensic samples on the EPIC methylation array (see *Methods*). Unfortunately, none of these passed post-assay quality control metrics and therefore could not be reliably analyzed alongside the other samples. It is likely that the DNA in these samples was already too degraded for quantitative measures of DNA methylation to be recovered after a few years of exposure to the elements and taphonomic processes. A common type of DNA damage occurs when cytosines are spontaneously degraded into thymines if they are methylated or uracils if they are unmethylated. Bisulphite treatment, which precedes most DNA methylation assay techniques, also converts unmethylated cytosines to uracils that are later read as thymines, but does not affect methylated cytosines; the attached methyl group ‘protects’ those cytosines from being mutated. DNA methylation levels are subsequently measured from the difference between the cytosines and thymines at a given CpG site. Therefore, post-mortem cytosine damage will generally make methylated cytosines appear unmethylated, a phenomenon that is the likely explanation for the large difference observed between the median methylated and unmethylated signals in the forensic samples (Figure 1).

The poor quality of our forensic DNA samples does not negate the utility of bone-specific DNA methylation-based age predictor. It does, however, highlight several practical considerations related to the application of any method that relies on good-quality DNA, including preservation and skeletal element choice . These issues have started to be investigated in the context of ancient DNA, where there is significant interest in understanding the DNA methylation profiles of individuals at different points in the past (Gokhman et al., 2017, 2014; Llamas et al., 2012; Pedersen et al., 2014; Seguin-Orlando et al., 2015; Smith et al., 2015). Studies have shown that it is possible to recover accurate measurements of DNA methylation at individual CpG site level even for samples that are thousands of years old (Llamas et al., 2012; Smith et al., 2015). However, the success of such bisulfite-based DNA methylation assays depends on the quality and preservation of the input DNA (Seguin-Orlando et al., 2015). Therefore, bone clock developed here may have substantial utility for forensic investigations when used in the appropriate contexts.

The forensic samples in the present study may have been from individuals that were particularly poorly preserved, but additionally, the skeletal element sampled, the intercondylar fossa of the femur, is not ideal for the recovery of DNA. Ancient DNA studies have shown that the petrous portion of the temporal bone is a good source of DNA that is relatively resistant to contamination and degradation (Gamba et al., 2014; Pinhasi et al., 2015). It is, however, difficult to access without significant destruction of the skull, which was permitted for the preserved samples but not for those sourced from the forensic collection. In ancient DNA studies, a sample’s collagen content is usually measured first before deciding if sequencing should be done, as this is a good indicator of preservation and DNA quality (Ovchinnikov et al., 2000). This may also be a worthwhile step to incorporate into a forensic investigation workflow when deciding which aging method is most appropriate for a given case.

In this study, we show that chronological age can be accurately estimated from bone-derived DNA from human adults. This model represents a promising molecular method that may be useful for building biological profiles of unknown individuals in forensic cases. Future research to focus on further refining the bone clock CpG set may allow for the development of more precise and, importantly, more cost-effective methods of aging individuals’ skeletal remains using DNA methylation markers.

## Acknowledgements

We would like to thank Daniel Wescott, Randall Susman, and Danny Soto for generously granting access to their respective collections of human remains. We would also like to thank Sjur Reppe for providing access to his bone DNA methylation dataset. Research funding and support for SG was provided by a National Institute of Justice Graduate Research Fellowship (2016-DN-BX-0011). JG was supported by a grant from the Undergraduate Research and Creative Activities summer program (URECA, Stony Brook University).

